# Ventral Tegmental Area dopamine projections to the hippocampus trigger long-term potentiation and contextual learning

**DOI:** 10.1101/2022.11.16.516747

**Authors:** Fares Sayegh, Catherine Macri, Juliana Pi Macedo, Camille Lejards, Claire Rampon, Laure Verret, Lionel Dahan

## Abstract

In most models, dopamine is not involved in the initiation but only in the long-term maintenance of LTP and memory. Conversely, we discovered that repeatedly stimulating Schaffer collaterals in concomitance with midbrain dopamine afferents to CA1 triggers hippocampal LTP. Moreover, we show that this dopamine pathway is involved in contextual learning. Thus, midbrain dopamine can play the role of a teaching signal triggering non-Hebbian LTP and allowing supervised learning.

## Manuscript

Hippocampal Long-Term Potentiation (LTP) is the main mechanism underlying episodic-like memories^1^. Classically, High Frequency Stimulation (HFS) was used to trigger LTP and elucidate the molecular mechanisms of its expression^2^. However, little is known about the molecular initiator of learning-induced LTP^3^.

Midbrain dopamine neurons fire in response to rewards^4^, aversive stimuli^5^ and salient events^6,7^, in other words, whenever something has to be learnt. They were thus proposed to provide a teaching signal to the hippocampus^8^. However, since dopamine antagonists spare the early phase of LTP while blocking the late maintenance of LTP requiring protein synthesis ^9,10^ and synaptic tagging^11^, research has mainly focused on its role as mediating novelty-induced enhancement of memory and behavioral tagging^12^. Although systemic injection of dopamine antagonists prevents learning-induced hippocampal plasticity^13^, no study addressed whether and how midbrain dopamine may trigger hippocampal LTP^14^, which is the missing part for fully understanding dopamine’s role in allowing supervised learning outside the Hebbian framework^15^.

Moreover, the source of hippocampal dopamine innervation is debated^7,16^. Indeed, while recent data show that stimulating midbrain dopamine projections to the hippocampus originating from the lateral Ventral Tegmental Area (VTA) are sufficient to facilitate contextual fear conditioning^16^, two earlier studies suggest that inputs arising from the locus coeruleus (LC) noradrenergic neurons constitute the main dopaminergic input to the hippocampus^7,17^.

Thus, we optogenetically manipulated VTA midbrain dopamine neurons and tested whether these inputs to the hippocampus trigger LTP. We injected a viral vector expressing the ultra-fast excitatory ChETA^18^ rhodopsin (AAV2-Ef1a-DIO-ChETA-EYFP), or its EYFP control (AAV2-Ef1a-DIO-EYFP), into the VTA of DAT::Cre^19^ mice. Both vectors induced an efficient and specific transfection of dopamine neurons in the VTA and medial Substantia Nigra pars compacta (Figure 1a). We performed electrophysiological recording of evoked field potentials at Schaeffer collaterals *in vivo* in urethane anesthetized mice. An optic fiber was placed into the glass recording pipette to phasically stimulate midbrain dopamine inputs to the recorded area in CA1 (Figure 1b).

**figure 1:**
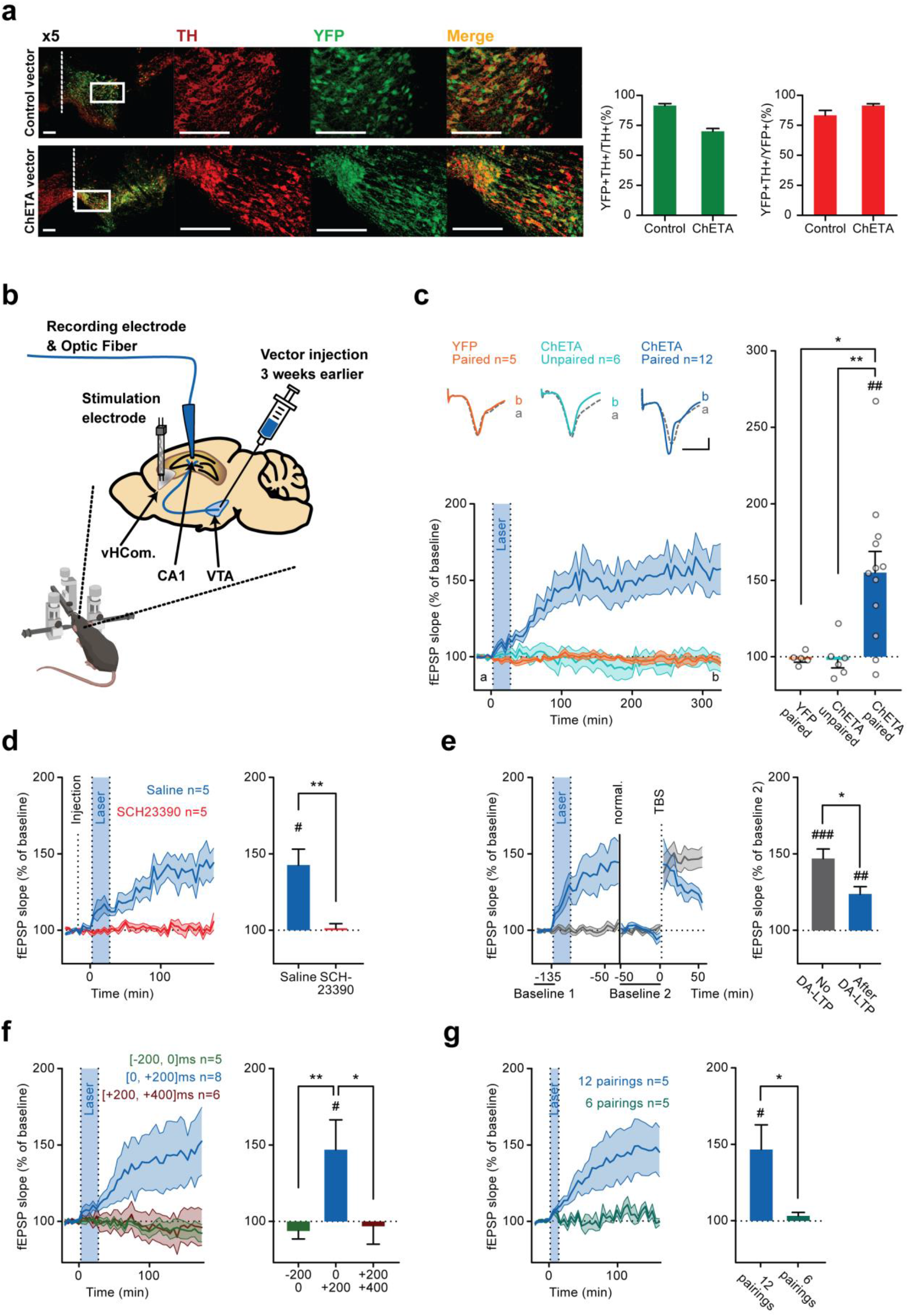
Midbrain dopamine triggers a long lasting, D1/5R dependent, increase of synaptic transmission in CA1, which occludes TBS-triggered LTP. **a**. Transfected brain slices were stained for Tyrosine Hydroxylase (TH) and YFP. Dashed line represents midline; rectangle represents sampled area captured at x20 zoom. Transfection and Specificity are represented as the number of double stained cells devised by TH+ cells and YFP+ cells, respectively; Transfection was 70±2% for mice transfected with ChETA and 91.5±2% for mice transfected with YFP, and Specificity 91.5±2% for mice transfected with ChETA and 83.4±4% for mice transfected with YFP. **b**. a schema representing the procedure. DAT::Cre mice were injected with either floxed YFP-coding vectors or ChETA and YFP-coding vectors in their Ventral Tegmental Area (VTA). Three weeks later, anesthetized in vivo electrophysiology recording of CA1 response to Schaffer collaterals electric stimulation in the ventral hippocampal commissure (vHCom.). Optic stimulation was delivered through the glass recording pipette. **c**. Effect of 50 coupling of dopamine axons photo-stimulations (4ms pulses at 50Hz during 400ms) to 50 electrical stimulations of Schaffer Collaterals (0,1ms, delivered 200ms after the onset of the light burst) (blue shaded part of the timeline). When stimulations were simultaneously coupled in ChETA injected mice, these couplings induced 55±14% increase in fEPSP slopes (Dark Blue, ChETA), we call this phenomenon DA-LTP. No such increase was observed when electrical and optogenetic stimulations were separated by 15 seconds (−2.1±5%, Light Blue, ChETA Unpaired), neither in mice injected with control vectors and simultaneously stimulated (−1.7±1.8%, Orange, YFP). **d**. SCH23390 injected 20 minutes prior to the coupling (dashed line); EPSP slope increase was no longer observed (+1.1±3% red, SCH23390). NaCl 0.9% injections did not affect the effect of the couplings (+42±10% Dark Blue, Saline). **e**. We Induced TBS LTP (dashed line, TBS) 90 minutes after the end of the couplings (and 50 minutes after renormalizing field slopes, full line) in 7 mice that received DA-LTP (Dark Blue, DA-LTP) and 7 mice that did not (Grey, No DA-LTP). Both groups, DA-LTP and No DA LTP, showed similar TBS induction of LTP. However, LTP in DA-LTP group degraded quickly (47.2±6% for No DA-LTP vs 23.9±5% DA-LTP). **f**. We used shorter Laser bursts (200ms) to determine the time window for DA-LTP induction. DA-LTP was induced when photo stimulations were delivered in the time window (0 to 200 ms) in relation to the electrical stimulation of SC (Dark Blue, +46±20%). No such increase was observed neither when photo stimulations were delivered (−200 to 0 ms) (Green, -6±5%) nor when photo stimulations were delivered (+200 to +400 ms) (Dark red, -3.2±12%) in relation to electrical stimulation of SC. **g**. 12 pairings of photo stimulations (0 to 200 ms in relation to SC electrical stimulations) were sufficient to induce DA-LTP (+46.5±16%, 12 pairings, Dark Blue), but not 6 (+3.3±2%). For panels **c-g**, Timelines of each group on the left, mean changes quantified by averaging the last 25 minutes of the recording for each mouse to the right. * p<0.05 Mann-Whitney, ** p<0.01 Mann-Whitney (after significant Kruskal Wallis). # p<0.05 t-test vs. 0. ## p<0.01 t-test vs. 0. ### p<0.001 t-test vs 0.

Our main finding is that fifty pairings of electrical stimulations of Schaffer collaterals delivered concomitantly with light bursts (burst duration: 400ms) induced an increase in the slope of Schaffer collaterals’ field potentials (+55±14%) (figure 1c, ChETA Paired). Interestingly, this synaptic plasticity developed progressively over 90 minutes and stayed stable for at least 5 hours after the last coupling. We called this phenomenon DA-LTP. Conversely, this protocol induced no plasticity in mice transfected with the control vector (figure 1c, YFP Paired) or in mice transfected with ChETA vector but receiving optogenetic stimulations unpaired with the electrical stimulation of the Schaffer collaterals (figure 1c, ChETA Unpaired).

DA-LTP was completely prevented by intraperitoneal injection of D1/5R antagonist SCH23390 (0.05mg/kg) which did not alter synaptic transmission (figure 1d). To test whether DA-LTP and classical LTP shared similar mechanisms, we performed an occlusion experiment. A train of theta burst stimulation (TBS) at the Schaffer collaterals induced a classical LTP of lower magnitude in mice that previously received a protocol inducing DA-LTP as compared to mice that did not (receiving ChETA Unpaired or YFP Paired protocols, figure 1e). Thus, DA-LTP partially occluded LTP triggered by TBS, supporting the hypothesis of a common mechanism of expression and maintenance for both forms of LTP.

In order to determine the time window of the pairing required to trigger DA-LTP, we reduced light bursts duration from 400ms to 200ms, which also better mimics natural burst of dopamine^20^. DA-LTP was triggered when dopamine terminals stimulation was delivered concomitantly with the stimulation of Schaffer collaterals and spanning for the next 200ms, but not when it started before or 200ms after the electrical stimulation (figure 1f). By progressively reducing the number of pairings, we found that 12 pairings were enough to induce a full DA-LTP, while 6 pairings had no lasting effect (figure 1g). Hence, triggering DA-LTP requires 7 to 12 dopamine stimulations mimicking naturally occurring bursts. Here we demonstrated that repeated bursts of activity of midbrain dopamine afferents to the hippocampus trigger LTP at coactivated Schaffer collaterals. Translated in natural conditions, this would mean that when dopamine cells innervating the hippocampus fire bursts of action potentials, the Schaeffer Collaterals synapses that are concomitantly activated – at a 200ms time scale– would be potentiated. In this regard, the requirement of 7 to 12 pairings to trigger LTP could allow selecting only the set of synapses that relates to unpredicted stimuli which had triggered the dopamine burst and was reliably concomitant. Interestingly, in regard to its slow developing time course, DA-LTP is similar to the potentiation triggered by bath application of D1/5 agonists^21,22^ or by aversive contextual learning^13,23^. These characteristics of midbrain dopamine inputs to the hippocampus correspond to the definition of the teaching signal previously hypothesized to trigger non-Hebbian LTP in response to rewarding, aversive or neutral unpredicted events^14,15^.

To test the implication of this pathway in hippocampus dependent learning, we studied contextual learning. Indeed, in earlier work we used a modified version of contextual fear conditioning called Context Pre-exposure Facilitation Effect (CPFE) to demonstrate that D1/5R are necessary for learning a new context in absence of any rewarding or aversive cue^24^. In this procedure, mice learn separately the context and its association with the electric shock on two consecutive days (figure 2a). We first validated this CPFE protocol in DAT::Cre mice. Control mice, not pre-exposed to the context, and mice pre-exposed for 30sec exhibited similar low levels of freezing (37±6% and 42.1±5%, respectively, figure 2b). Conversely, 2 or 8 min of pre-exposure allowed contextual learning, as revealed by similar increase in freezing behavior during the test (62.7±4% and 61.7±5%, figure 2b). We then manipulated this pathway during the pre-exposure session. On the one hand, mice expressing ChETA in VTA dopamine neurons and receiving 90 bursts of light bilaterally in the dorsal hippocampus all along 30 seconds of pre-exposure exhibited high levels of freezing (62.8±4%), significantly higher than those observed in YFP mice (45.6±3%; figure 2d). On the other hand, mice expressing the inhibitory opsin eNpHR3.0 in dopamine VTA neurons and pre-exposed to the context during 2 min while receiving a continuous green light illumination of the hippocampus showed low levels of freezing (47.2±6%), which are comparable to non-pre-exposed mice and significantly lower that of mice receiving the same treatment but injected with the YFP control vector (79.2±6%). In both experiments, freezing in an alternative context was considerably lower than in the conditioned context, and was not modified by dopaminergic activation or inhibition during pre-exposure (Figures 2d and 2e). Thus, midbrain dopamine inputs to the dorsal hippocampus are involved in setting up the cognitive representation of a novel context, so that it can be associated, the next day, with the electric shock. This corroborates our hypothesis stating that midbrain dopamine inputs to the dorsal hippocampus provide a signal triggering learning regardless of value inputs.

**figure 2:**
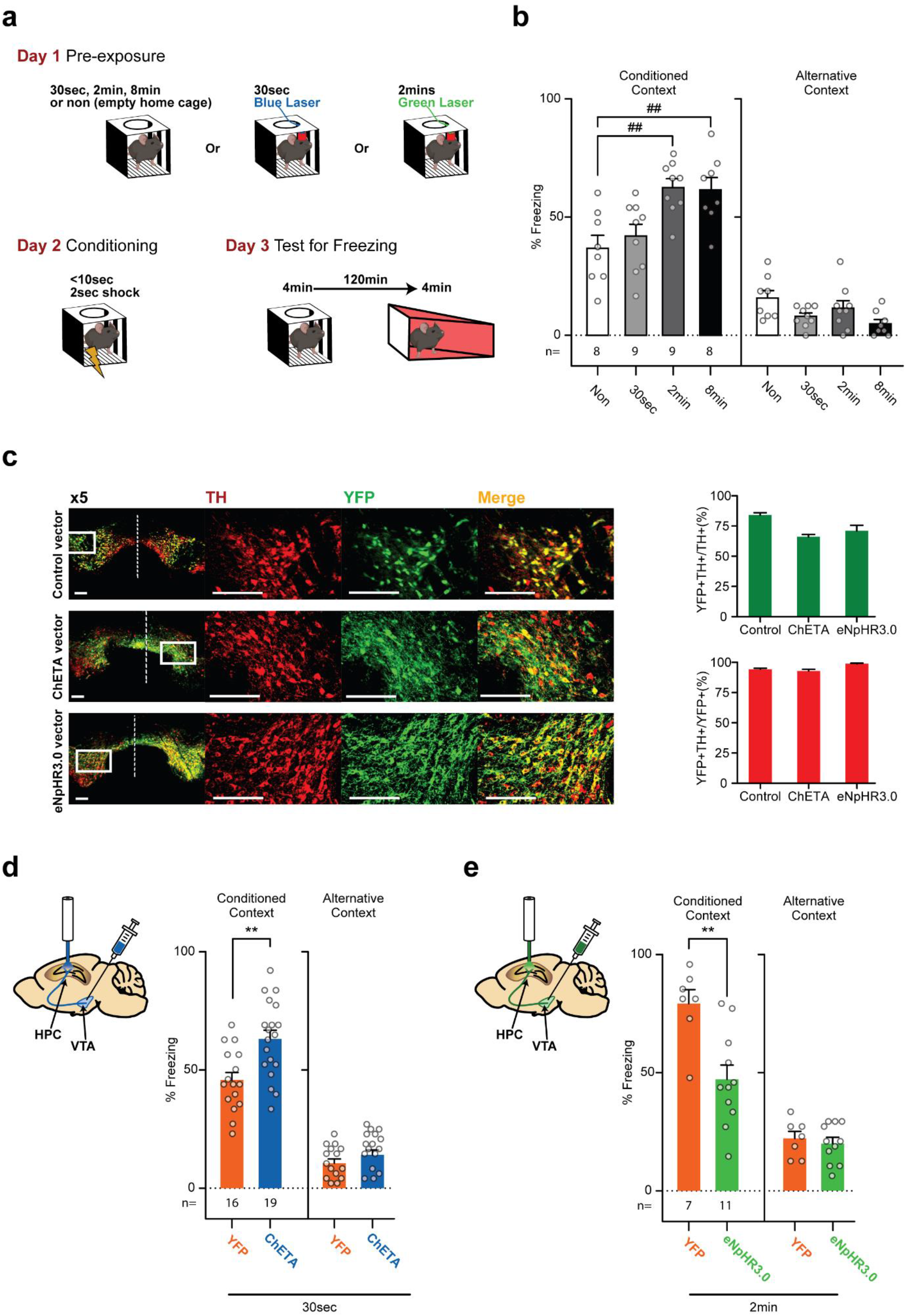
Midbrain dopamine in the hippocampus is responsible for learning a new context. **a**. a schema representing the procedure. DAT::Cre mice were used in a context pre-exposure facilitation effect paradigm. First, the effect of different pre-exposure durations was studied, and then the effect of optogenetic manipulation of midbrain dopamine afferents to the hippocampus was studied. On day 2 mice received and immediate shock and on day 3 freezing was tested in the conditioned context and in an alternative one. **b**. DAT::Cre mice with a 30sec pre-exposure did not freeze more than control non-pre-exposed group (42.1±5% vs 37±5%). 2min pre-exposure was sufficient to trigger a significant increase reaching levels comparable to those seen in the group with 8min pre-exposure (62.7±4% and 61.7±5%, respectively). **c**. Transfected brain slices were stained for Tyrosine Hydroxylase (TH) and YFP. Dashed line represents midline; rectangle represents sampled area captured at x20 zoom. Transfection and Specificity are represented as the number of double stained cells devised by TH+ cells and YFP+ cells, respectively; Transfection was 65.9±2% for mice transfected with ChETA, 83.9±2% for mice transfected with YFP and 70.8±5% for mice transfected with eNpHR3.0 and Specificity 93.3±2% for mice transfected with ChETA, 94.7±1% for mice transfected with YFP and 99.4% for mice transfected with eNpHR3.0. **d**. Mice were pre-exposed for 30 seconds on day one during which they received 90 bursts (burst duration: 200ms, 4ms pulses, @50Hz) of blue light (473nm, 10mW) bilaterally in the dorsal hippocampus. Freezing in the conditioned context increased in the ChETA-injected mice (62.8±4%) compared the YFP-injected (45.6±3%). Freezing in the alternative context was considerably lower than in the conditioned context and was not significantly changed due to dopaminergic activation during pre-exposure (10.6±2% for YFP and 14±2% for ChETA). **e**. Mice were pre-exposed for 2 minutes on day one during which they received continuous green light (532 nm, 10mW, starting 20s before placing the mouse in the context). Freezing in the conditioned context levels observed during the test on day 3 were lower for eNpHR3.0 vector injected mice (47.2±6%) in comparison to mice with control injection (YFP) (79.2±6%). Freezing in the alternative context was considerably lower than in the conditioned context and was not significantly changed due to dopaminergic inhibition’ during pre-exposure (19.9±6% for YFP and 22±3% for eNpHR3.0). ## p<0.01 Multiple comparisons following one-way ANOVA (p<0.001). ** p<0.01 t-test.

These results are in line with recent data showing that midbrain dopamine inputs to the hippocampus facilitate aversive memories^16^ and further show that this pathway is involved in learning a context in the absence of any association to aversive or rewarding events. Although this may seem inconsistent with previous work from Takeuchi *et al*. showing that dopamine released from LC, and not VTA, facilitates the long-term maintenance of fading memories, it is important to keep in mind that we tested if dopamine acted as a teaching signal by stimulating dopamine afferents during the learning episode. In contrast, Takeuchi *et al*. did so 30 minutes thereafter, in order to test for the role of dopamine in behavioral tagging^7^. Noteworthy, they also found a slowly developing increase in fEPSP of Schaeffer collaterals when strongly stimulating LC neurons in hippocampal slices. This is similar to our DA-LTP, although they did not test for the requirement of coincident stimulation, neither did they test if it would last longer than 40 minutes. In the same line, Kempadoo *et al*. showed that manipulating dopamine input from the LC enhanced learning in the object location task which does not involve a reward either^17^. Given they also show that most of the dopaminergic innervation to the dorsal hippocampus comes from the LC, they finally argued that LC has a greater influence on the dorsal hippocampus than on the VTA. However, a very recent study shows that LC inputs play a role in updating recently acquired contextual memories, but not in contextual learning per se^25^. Since we clearly show that midbrain projections influences learning very significantly, pathways arising either from LC or from midbrain monoaminergic neurons may play complimentary roles in both synaptic plasticity and behavior.

In summary, we show that midbrain dopamine projections to the hippocampus trigger DA-LTP though D1/5 receptors when dopamine is released repeatedly and concomitantly with Schaeffer collaterals activation. Furthermore, using context pre-exposure facilitation effect of contextual fear conditioning, we show that stimulating midbrain dopamine afferents to the hippocampus promotes contextual learning, while their inhibition hinders it and propose that dopamine, through the triggering of hippocampal LTP, may act as a teaching signal, selecting relevant inputs to be memorized.

## References

1. Morris, R.G.M.M. Philos. Trans. R. Soc. B Biol. Sci. 358, 643–647 (2003).

2. Baltaci, S.B., Mogulkoc, R. & Baltaci, A.K. Neurochem. Res. 44, 281–296 (2019).

3. Whitlock, J.R., Heynen, A.J., Shuler, M.G. & Bear, M.F. Science (80-.). 313, 1093–1097 (2006).

4. Ljungberg, T., Apicella, P. & Schultz, W. J. Neurophysiol. 67, 145–163 (1992).

5. Matsumoto, M. & Hikosaka, O. Nature 459, 837–841 (2009).

6. Schultz, W. Adv. Pharmacol. 42, 686–690 (1997).

7. Takeuchi, T. et al. Nature 537, 357–362 (2016).

8. Harley, C.W. Neural Plast. 11, 191–204 (2004).

9. Lemon, N. & Manahan-Vaughan, D. J. Neurosci. 26, 7723–7729 (2006).

10. Frey, U., Huang, Y.-Y. & Kandel, E.R. Science (80-.). 260, 1661–1664 (1993).

11. Sajikumar, S. & Frey, J.U. Neurobiol. Learn. Mem. 82, 12–25 (2004).

12. Rossato, J.I., Bevilaqua, L.R.M., Izquierdo, I., Medina, J.H. & Cammarota, M. Science (80-.).325, 1017–1020 (2009).

13. Broussard, J.I. et al. Cell Rep. 14, 1930–1939 (2016).

14. Otmakhova, N., Duzel, E., Deutch, A.Y. & Lisman, J. Intrinsically Motiv. Learn. Nat. Artif. Syst. 235–254 (2013).doi:10.1007/978-3-642-32375-1_10

15. Gerstner, W., Lehmann, M., Liakoni, V., Corneil, D. & Brea, J. Front. Neural Circuits 12, 1–16 (2018).

16. Tsetsenis, T. et al. Proc. Natl. Acad. Sci. 118, e2111069118 (2021).

17. Kempadoo, K.A., Mosharov, E. V., Choi, S.J., Sulzer, D. & Kandel, E.R. Proc. Natl. Acad. Sci. U. S. A. 113, 14835–14840 (2016).

18. Gunaydin, L.A. et al. Nat. Neurosci. 13, 387–392 (2010).

19. Turiault, M. et al. FEBS J. 274, 3568–3577 (2007).

20. Dahan, L. et al. Neuropsychopharmacology 32, 1232–1241 (2007).

21. Sajikumar, S., Navakkode, S. & Frey, J.U. Learn. Mem. 15, 46–49 (2008).

22. Navakkode, S., Sajikumar, S. & Frey, J.U. Neuropharmacology 52, 1547–1554 (2007).

23. Subramaniyan, M. et al. Hippocampus 31, 1154–1175 (2021).

24. Sayegh, F. et al. Learn. Mem. 29, 142–145 (2022).

25. Chowdhury, A. et al. Neuron 1–15 (2022).doi:10.1016/j.neuron.2022.08.001

