## Supplementary Methods for "Ventral Tegmental Area dopamine projections to the hippocampus trigger long-term potentiation and contextual learning"

**Animals**

Mice 2-6.5 month old male DAT::Cre[19] mice were bred in-house. DAT::Cre mice are descendants of FVB/N with Cre-recombinase under the control of the dopamine transporter promoter (DAT) inserted into a bacterial artificial chromosome (BAC) then mice were backcrossed more than 15 times on the C57BL/6J line. These animals are placed in stalls in groups of 3 to 5 individuals per cage, with food and water available ad libitum, under a 12h/12h day/night cycle (day from 8h to 20h) at 22 ± 1 °C. All experiments were performed in strict accordance with the recommendations of the European Union (86/609/EEC) and the French National Committee (87/848).

**Electrophysiology**

Anesthesia was induced using Isoflurane then maintained with an injection of urethane (1.5±0.2 mg/g) during the recording period. In the stereotaxic apparatus mice received an incision and a craniotomy, then stimulating electrodes were placed at the ventral hippocampal commissure (Coordinates: -0.3mm AP, -0.5mm ML and 2.3 DV) to evoke field excitatory postsynaptic potential (fEPSP) recorded at the stratum radiatum of area CA1 recorded using a glass micro-pipette (NaCl 2M filled 4microns thick 0.8-1.1 Megaohm @100Hz). Recordings were performed using an A-M Systems model 1800 amplifier (gain = 100, bandwidth of 0.1Hz to 10kHz) and stimulations using 2100 Isolated Pulse Stimulator from A-M Systems both connected to CED micro 1401 which allowed sampling at 10kHz after 50Hz filtration using Humbug 50Hz Noise Eliminator, QuestScientific. Signals were visualized and analyzed using Spike2. The raw electrophysiological signal was first analyzed by averaging fEPSP waveforms every 5 minutes (10 fEPSP). The slope of the initial phase of each mean fEPSP is measured. A baseline was obtained by stimulating with a current providing 70% of the maximal response recorded (0.05-1mA biphasic 100µs stimulations). We considered 25min of recording (i.e. five mean fEPSPs) with a slope between 95 and 105% of the average as a stable baseline and the evolution of the fEPSP slope should not follow a linear change. We then delivered the coupling of glutamatergic stimulation and dopaminergic optic stimulation after which, a follow up took place. For the occlusion experiment, mice received a HFS protocol of LTP. This protocol followed Theta Burst Stimulation (TBS) pattern, for which we used 4 trains (30s inter-train interval) of 4 bursts (200ms inter-burst interval) of 5 stimulations at 100Hz. At the end of the recording, the glass pipette is replaced with one filled with Chicago blue 2% in acetate buffer 0.5M. We searched for the same recorded fEPSP and the recording site was marked using electric expulsion of the dye with negative current of 20 µA, cycles 10 sec "on" / 10 sec "off" for 10 minutes. Finally, the mice were euthanized with a lethal injection of pentobarbital, and then, received an intracardial infusion of 0.9% NaCl solution and the brains were collected for immunohistochemical verification.

**Vectors**

For optogenetics manipulations we used UNC Vector Core provided vectors; AAV2-Ef1a-DIO-ChETA-EYFP, AAV2-Ef1a-DIO-EYFP or AAV2-Ef1a-DIO-eNpHR3.0-EYFP at original concentrations at purchase (3.5-5x1012 molecule/mL). For electrophysiological experiments, mice received 2 unilateral injections of 0.5µL/Site at (-3.3mm AP, +0.6mm ML and -4.0/-4.6mm DV). For behavioral experiments mice received 2 bilateral injections 0.3µL/Site at (-3.3mm AP, ±0.6mm ML and -4.0/-4.6mm DV).

**Laser delivery**

Two lasers were used for this work 473 nm DPSS Laserglow (blue light for ChETA activation) and 532 nm DPSS Laserglow (green light for eNpHR3.0 activation). Light was then connected to the head of the mouse either through 200 µm patch chords, then through 200 µm nude optic fibers and finally, through the implanted cannulas (for behavioral studies). For electrophysiology, longer fiber optic cannulas were placed inside the recording glass micro-pipette that led the light to the recording site. The intensity was set to 10 mW at the implantable tip. For inhibition, mice with eNpHR3.0 vectors received one continuous pulse (140 seconds long) covering 20 seconds before context pre-exposure the 2 minutes of pre-exposure. For stimulation, mice received either 200ms or 400ms bursts (4ms pulses, 50Hz) depending on the protocol. These pulses were driven using Model 2100 Isolated Pulse Stimulator from A-M Systems and Spike2 connected through Micro1401 from CED.

**Behavioral task**

Context Pre-exposure Facilitation Effect (CPFE) was performed as described in a former paper[24], on day 1, mice were allowed to explore the context (27x27x27cm, with horizontal black and white stripes on a wall and vertical ones on the opposite, metal rods floor, white light) during 30 seconds, 2 minutes or 8 minutes. We used as control a non-pre-exposed group that explored an empty home cage for 8 minutes. On day 2, all animals received a very brief conditioning session (<10 seconds) during which they received one electric shock (0.7 mA, 2s) in the conditioning context. Tests took place on day 3; freezing behavior was measured in the same context for 4 minutes to assess contextual memory. An hour and a half later, freezing was measured in an alternative context (triangular shape, transparent Plexiglas walls and floor, red light) for 4 minutes to assess generalized fear, mice showing >33.33% generalization were excluded from the analysis (1 from non-pre-exposed mice and 4 YFP). Freezing was scored each 5 seconds by two independent experimenters blind to the experimental condition and expressed as a percentage of the sampled time spent freezing. In order to respect parametric requirements for ANOVA tests, percentages (P) were transformed using the equation $Q = \mathrm{Asin}(\sqrt{P}/100)$[24]. To make sure that odor association did not happen, we used nonalcoholic makeup remover wipes (Ysiance, aloe vera) to clean the contexts on the first two days and alcohol solution on day three. The experimenter who conducted the conditioning session on day 2 was different from the experimenter on days 1 and 3 in order to avoid any association between the context, the experimenter and the shock. Before optogenetic manipulations, mice were first habituated to the connection of the optic fibers during the week preceding the experiments by thrice connecting them to patch cords and allowing them to explore an empty home cage once every two days. Mice injected with either ChETA coding vector or its control were pre-exposed to the context for 30 seconds during which they received 90 bursts of blue light bilaterally in the dorsal hippocampus (burst duration: 200ms, 4ms pulses @50Hz, 473nm, 10mW). Two other groups, injected with either eNpHR3.0 coding vector or its control, and were pre-exposed to the context for 2 minutes during which they received a continuous green light bilateral illumination of the hippocampus (532nm, 10mW).

**Immunohistochemistry**

To measure the efficiency of the transfection, we carried out a double immunohistochemical labeling revealing the reporter protein eYFP, and Tyrosine Hydroxylase (TH) the enzyme necessary for dopamine production, to visualize the transfected cells and dopaminergic cells, respectively. The animals were anesthetized (pentobarbital) before performing an intracardiac infusion with 0.9% NaCl solution (20-30s, 20mL/min). The brains were then removed and placed in 4% PFA solution for 24-72hr, then rinsed with 0.1M PBS. Finally, the brains were stored in a 30% sucrose solution containing 0.1% azide. These brains were later sectioned into several serial 40µm sections with a freezer microtome and then stored in a cryoprotectant solution. For immunohistochemistry staining, on day one, sections were washed in 0.1M phosphate buffered saline containing 0.25% triton (PBST), placed for 15min in a solution containing 10% H2O2 and 10% methanol (in PBST), to block the endogenous peroxidase. Then two rinses of 10 min with PBST took place, after that sections were placed for 1 hour in a solution saturating non-specific bonds (BlockNsp: 5% donkey serum in PBST). Finally, they were incubated overnight at room temperature in a solution of BlockNsp containing the primary antibodies (goat anti-YFP 1:2500 (Rockland, 600101215) and rabbit anti-TH 1:1000 (Millipore, AB152)). The next day, the sections were twice rinsed in PBST before being placed for 1 hour and a half in a solution containing the fluorescent secondary antibodies (donkey anti-goat A488 1:250 (Thermofisher, A11055) and donkey anti-rabbit A555 1:250 (Thermofisher, A31572)). Finally, the sections were twice rinsed in PBST before mounting them on slides. Slide covers were glued with Mowiol containing Hoechst (1: 10,000) in order to mark the nuclei. Once dry, the slides were observed using a Leica fluorescence microscope, a sampled transfection zone was counted for each mouse (using Mercator software). Sections were photographed using the same software and these photos were retouched using ImageJ. Slides were then stored at 4 ° C. Transfection analysis relied on two metrics, specificity and efficiency by studying the collocation of TH expressing cells and YFP expressing cells.

Specificity was calculated by the equation:

$$specificity\% = \frac{\#[\mathrm{TH+, YFP+}]}{\#[\mathrm{YFP+}]} * 100$$

Efficiency was calculated by the equation:

$$transfection\% = \frac{\#[\mathrm{TH+, YFP+}]}{\#[\mathrm{TH+}]} * 100$$

**Data analysis**

Number of animals per group was calculated using minitab and aiming at 90% statistical power. All statistical tests and figures were done using GraphPad Prism 8 and graphs were retouched using Adobe Illustrator CS6.
